# Systematic Transmission Electron Microscopy-Based Identification and 3D Reconstruction of Cellular Degradation Machinery

**DOI:** 10.1101/2021.09.26.461841

**Authors:** Kit Neikirk, Zer Vue, Prasanna Katti, Ben I. Rodriguez, Salem A. Omer, Jianqiang Shao, Trace Christensen, Edgar Garza Lopez, Andrea Marshall, Caroline B. Palavicino-Maggio, Jessica Ponce, Ahmad Alghanem, Larry Vang, Taylor Barongan, Heather K. Beasley, Taylor Rodman, Margaret Mungai, Marcelo Correia, Vernat Exil, Sandra A. Murray, Jeffrey L. Salisbury, Brian Glancy, Renata O. Pereira, E. Dale Abel, Antentor O. Hinton

## Abstract

Many interconnected degradation machineries including autophagosomes, lysosomes, and endosomes work in tandem to conduct autophagy, an intracellular degradation system that is crucial for cellular homeostasis. Altered autophagy contributes to the pathophysiology of various diseases, including cancers and metabolic diseases. Although many studies have investigated autophagy to elucidate disease pathogenesis, identification of specific components of the autophagy machinery has been challenging. The goal of this paper is to describe an approach to reproducibly identify and distinguish subcellular structures involved in macro autophagy. We provide methods that help avoid common pitfalls, including a detailed explanation for distinguishing lysosomes and lipid droplets and discuss differences between autophagosomes and inclusion bodies. These methods are based on using transmission electron microscopy (TEM), capable of generating nanometer-scale micrographs of cellular degradation components in a fixed sample. We also utilize serial block face-scanning electron microscopy (SBF-SEM) to offer a protocol for visualizing 3D morphology of degradation machinery. In addition to TEM and 3D reconstruction, we discuss other imaging techniques, such as immunofluorescence and immunogold labeling that can be utilized to reliably and accurately classify cellular organelles. Our results show how these methods may be used to accurately quantify the cellular degradation machinery under various conditions, such as treatment with the endoplasmic reticulum stressor thapsigargin or ablation of the dynamin-related protein 1.

## 1 Introduction

Macroautophagy, the mechanism by which intracellular components or damaged organelles are removed and degraded to maintain cellular homeostasis [1], has much relevance in the fields of disease research and drug development. Although poorly understood, autophagy regulation is broadly implicated in disease pathogenesis, with both overactive and underactive autophagy having negative consequences, including malignant transformation and cellular proliferation in cancer or accumulation of ineffective cells in neurodegenerative diseases [2,3]. Autophagic processes differ depending on their activation pathways, being either non-selective or selective for specific cellular organelles or proteins [1,2]. Growing interest in neurodegenerative and other diseases with autophagy implications has highlighted its consequential roles in key biological processes [1,4].

The complex, regulated macro autophagic process involves structures that also contribute to the cellular recycling machinery specifically, autophagosomes and lysosomes (Figure 1A). The main stages of autophagy include initiation, elongation, autophagosome formation, autophagosome recruitment and maturation, fusion, and degradation [1–3]. In the initiation stage, often triggered by amino acid starvation, sack-like autophagosome precursors, called phagophores, assemble adjacently to the endoplasmic reticulum, typically at mitochondria-associated endoplasmic reticulum (ER) membrane (MAM) (Figure 1B) [5]. The phagophore resides proximal to the ER as an empty, unclosed membrane. As materials are delivered to the phagophore, the membrane closes to seal the organelle, transforming the phagophore into an autophagosome, which carries cytoplasmic components, cargo proteins or organelles designated for degradation. In many cases, the autophagosome then ultimately fuses with a lysosome, containing hydrolases and permeases, to form an autolysosome and initiate degradation (Figure 1B). In some cases, the autophagosome may mature through an intermediate prior to lysosome fusion known as an amphisome [6]. Importantly, this endosome fusion event may occur to allow for retrograde transport motility, so amphisomes can move to lysosome dense areas [6,7]. In either case, after autolysosome formation, the resulting macromolecules are released through permeases and the ultimate fate of autolysosomes following this process is still unclear [8]. Some autolysosome components can be used to reform lysosomes or become part of new phagophore membranes. Macromolecules released into the cytosol are recycled for use in other biological functions [1–3,9].

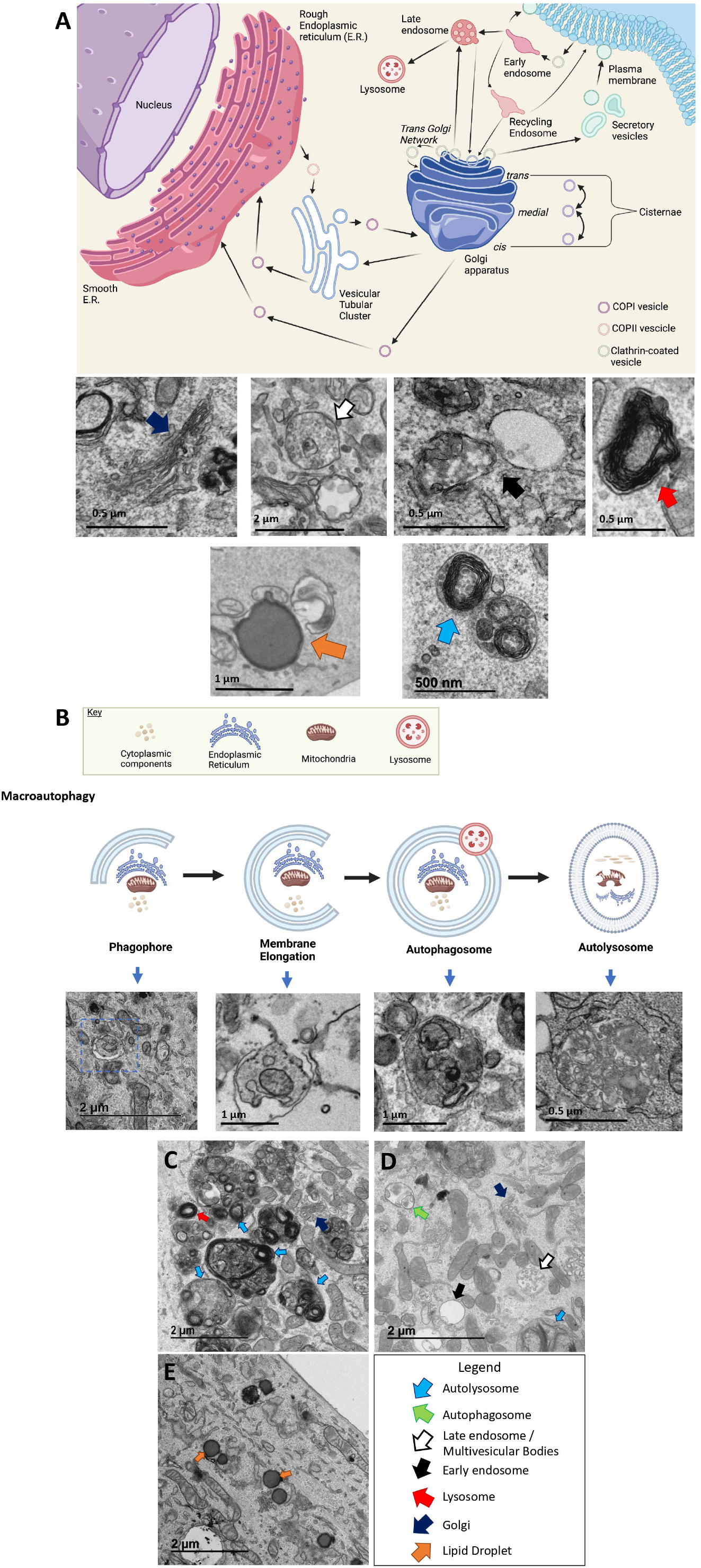

Transmission electron microscopy (TEM) has advanced autophagy research by enabling the study of subcellular components at high resolution [4]. By transmitting electrons through ultrathin sections of fixed and embedded samples, TEM generates nanometer-scale micrographs that allow study of autophagic processes [10–12]. For example, past studies have utilized TEM to show autophagosome formation at MAMs and have delineated the maturation process described above [10,12]. Our method uses the free, open-source ImageJ software platform to analyze TEM micrographs of autophagic components [13], enabling image quantitation and statistical analysis [11]. Here we applied this established TEM method recently described by Lam et al. [11] to analyze mitochondria and ER.

While TEM is powerful for identification, it only allows for two-dimensional (2D) visualization of organelles. This may not always be an accurate representation since organelles are three-dimensional (3D) objects. Therefore, we also included reconstrucions from serial block face-scanning electron microscopy (SBF-SEM) [14], which functions by slicing a sample in the z-axis to obtain orthoslices [15]. These orthoslices can then be hand-segmented, one-by-one, to allow for the 3D rendereings of an organelle to be evaluated [16]. While this allows for more accurate representation, there is a much larger time and cost associated with it than TEM quantification. Here we also utilized an established SBF-SEM method recently described by Garza-Lopez et al. [16], to perform 3D reconstruction of recycling machinery organelle alongside TEM quantifications.

The success of our modified protocol depends on proper identification of cellular degradation components. In Figure 1, we showcase how typical organelles should appear conventionally: lysosomes (Figure 1C, red arrows), autolysosomes (Figure 1C-D, blue arrows), golgi (Figure 1C-D, dark blue arrow), autophagosome (Figure 1D, green arrow), early endosome (Figure 1D, black arrow), late endosome (Figure 1D, black outlined white arrow) and lipid droplets (Figure 1E, orange arrow). However, identification is complicated by diverse morphologies with recycling machinery sometimes not presenting typical presentation. For example, lysosomes are usually depicted as spherical, typically ranging from 0.2 to 0.5 micrometers in diameter, although they can also commonly present from 0.05 to 1 micrometers (Figure 2A–D; Table 1) [17–19]. Lysosomes are typically transported by microtubules to the region around the microtubule-organizing center [20]; however, intracellular conditions, such as raised pH, cause lysosomes to migrate toward the cell membrane [21]. Lysosomes are classified as primary, secondary, or tertiary, depending on their digestive activities and their formation process. Lysosome identification is further complicated by their tendency to feature multiple membranes [22]. In quantifying lysosomes, researchers must avoid misidentifying them as multilamellar vesicles, also identified as late endosomes or multilamellar bodies (MLBs; Figure 2E–F; Table 1), which contain lipids within a central compartment surrounded by many membrane bilayers [23]. Lysosomes should not be mistaken for multi-inclusion bodies and multivesicular bodies (MVBs), also known as pre-vacuolar compartments, that are an intermediary structure between vacuoles and the *trans*-Golgi network (Table 1) [24]. Although these structures are related to autophagy, they differ from lysosomes and autophagosomes and should be excluded from both their quantifications [25]. Lysosomal membranes have highly organized inner folds and lysosomal enzymes have a distinct, darker, and more consistent appearance than the lipids found in MLBs (Figure 2A–F; Table 1). MLB lipids can also appear as dots that speckle the MLB interior, which can be used to differentiate MLBs from lysosomes, although these dots can also be mistaken for cargo in secondary lysosomes (Figure 1A-C; Table 1). Despite these considerations, accurate and reproducible lysosome identification based solely on TEM imaging may be inconsistent, necessitating use of imaging techniques, such as fluorescent staining. Not only do we need to positively identify lysosomes, but autophagosomes must be identified, characterized, and distinguished from lysosomes and other structures.

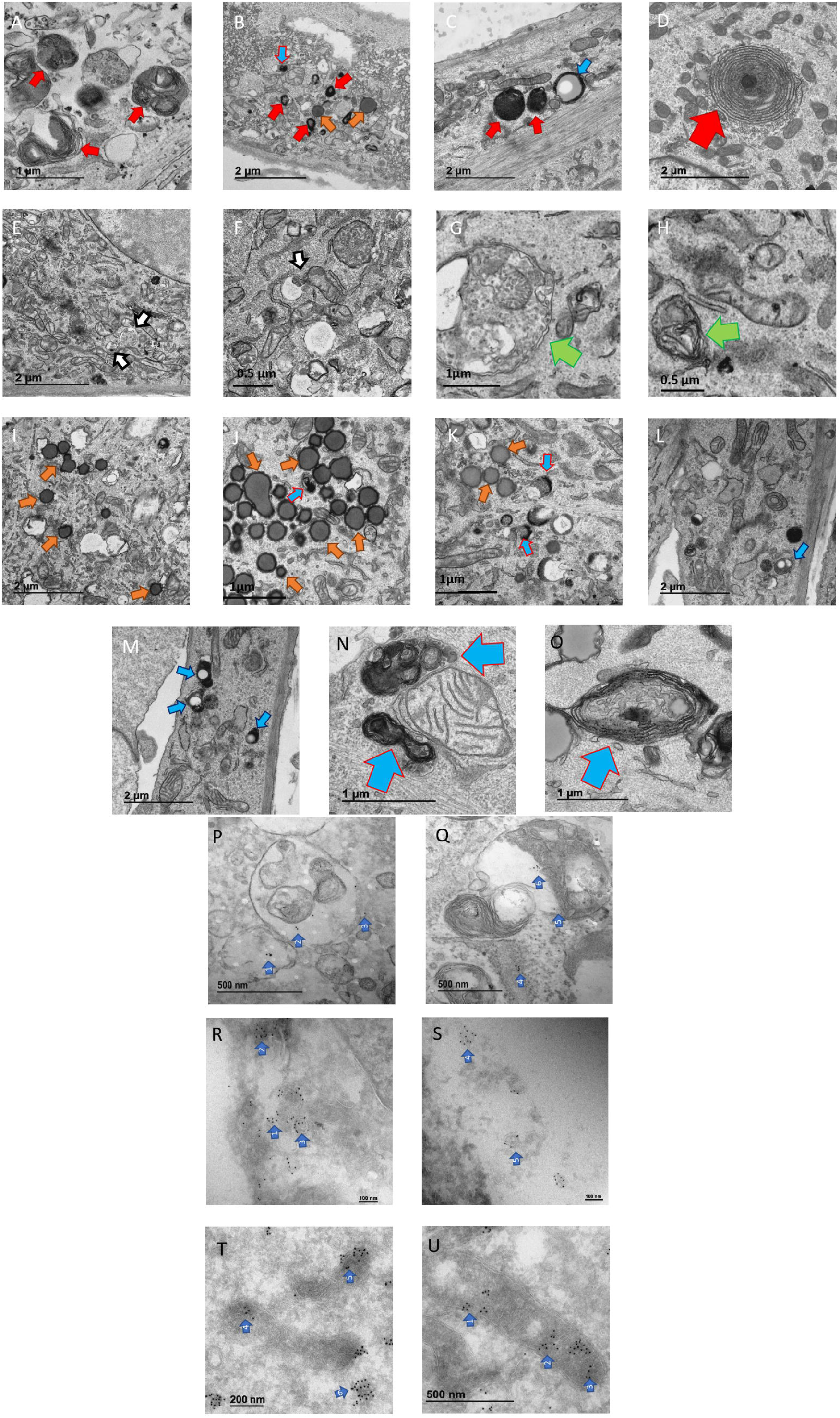

Autophagosome appearance can also vary depending on the cargo, further complicating their identification (Figure 2G-H; Table 1). Autophagosomes typically have clear double-limiting membranes that appear darker than the rest of the TEM image, separated by a small electron lucent space. However, autophagosomes may also appear with a single membrane or with several separate membranes due to fixation techniques (Figure 2G-H; Table 1) [26]. These diverse membrane presentations can cause misidentification of malformed mitochondria or ring-shaped ER as autophagosomes. Because MVBs and MLBs only display a single membrane (Figure 2E-F; Table 1), presence of a second membrane, in addition to inner recycled ribosomes, more circular shapes representing cargo, or more internal lipids, all characterize autophagosomes (Figure 2G-H). [23,26]. However, improper fixation techniques may cause one of autophagosome limiting membranes to be invisible [26]. MVBs are also generally smaller than autophagosomes and lysosomes, whereas MLBs are larger, sometimes up to ten times the size of typical lysosomes [17,23]. During identification, certain autophagosome types can be included or excluded. For example, if structures appear empty or lacking material, they are likely not involved in degradation and may be excluded. For example, in some disease states, autophagosomes may dysfunction in cargo recruitment and are known as “empty” autophagosomes [27]. Therefore, a key trait of autophagosomes is the presence of cargo. However, it is important to avoid classifying empty autophagosomes as lipid droplets (LDs; Figure 2I–K; Table 1). Similarly, if the body of a phagophore that has not yet closed (Figure 1B) to form an autophagosome, it should not be mistaken as an autophagosome [4,26]. Autophagosomes that are in the process of fusing with a lysosome, also known as lipofuscin granules (Figure 2C, 2E-D), but are not yet considered autolysosomes, can be classified as autophagosome-lysosome fusions, and be considered autolysosomes if their inclusion is consistent [1,2,28]. Since they, physiologically, are similar to autolysosomes, our criteria includes them in autolysosome quantifications [29]. These intermediate structures are identified by their much larger appearance, and contents of the lysosome and the autophagosome often appear to be interacting (Figure 2C, 2E-D).

Autophagosomes that contain limited cargo volumes (Table 1) can mimic large, irregularly-shaped lysosomes (Figure 2A; Table 1) [1,17]. Although autophagosomes can mainly be identified by two limiting membranes, because lipids are not reliably preserved during sample preparation, limiting membranes may vary in appearance, furthering potential for confusion [26]. Another key to identifying autophagosomes formed from non-selective phagophores is recognizing nearby cytoplasmic content inside of them; often autophagosomes can be identified by presenting cargo originating from cytoplasmic content nearby the organelle [30,31]. Furthermore, autolysosomes can differ in their presentation (Figure 2B-C, 2J-O), especially lipofuscin granule autolysosomes (Figure 2C, E-D; Table 1) which may be mistaken for LDs (Figure 2I-J; Table 1) or lysosomes (Figure 2A– D; Table 1). Careful consideration of these features is essential to properly identify these organelles.

Ultrastructural characteristics of organelles can be used to distinguish them from autophagic components. For example, the presence of regularly-spaced ribosomes, which typically appear as small black dots, or a thinner width wrapped around organelles such as mitochondria, strongly indicate that the structure is ER. Similarly, evidence of mitochondrial cristae or inner membrane folds can be used to identify mitochondrial structures. However, exact identification of specific organelles can be complicated. For example, partially degraded ribosomes can aggregate in autophagosomes, forming electron dense clumps as they degrade [26]. However, the presence of ribosomes alone does not confirm an object is an autophagosome, as it is possible to mistake circular cisterns of rough ER for autophagosomes [26]. Autophagosomes may be distinguished by their more circular presentation and higher-degree of clumped ribosomes [12,26].

Although basic processes of autophagosome formation are understood, specific pathways and degradation machinery require further study, such as lipid mobilization from LDs to provide energy. Like all organelles, LDs are targeted by autophagy for recycling and interestingly, macromolecules released by autophagy can be stored in new LDs, even under starvation conditions [32]. Thus, a consequence of autophagy is increased LDs within a cell. These lipid droplets can protect against ER stress and may protect against mitochondrial autophagy, known as mitophagy, by forming close mitochondria-to-lipid contacts [33]. Autophagy impacts nearly every cellular organelle due to its role in organelle degradation. For example, ER stress can initiate autophagy to recycle damaged ER membranes, contributing to healthy ER [34]. Similarly, impaired mitochondrial fission or other l dysfunctions due, for example, to impaired function of critical regulatory proteins, such as mitofusin 1 (MFN1), can trigger mitophagy to clear ineffective mitochondria [35,36]. Because all organelles interact with the cellular degradation machinery, understanding the dynamics between primary recycling organelles—lysosomes, autophagosomes, and autolysosomes—and other organelles is critical to fully appreciate the contributions and drivers of autophagy. Careful and accurate identification of components of the autophagy machinery are needed to advance our understanding of therapeutic effectiveness, whose mechanisms of action may involve autophagy. This effort may elucidate additional pathways that induce autophagy and clarify how autophagy contributes to disease prevention and progression [37]. Accurate characterization and quantitation of autophagy components requires proper identification of these degradation organelles and other subcellular structures involved in autophagic processes.

Although many studies have identified components of autophagosome and lysosome machinery, developing next-generation methods to rigorously identify and quantify these organelles is essential to establish standardized protocols, allowing data comparison [10,26,38–40]. Many of these structures are similar, but can differ from cell to cell and are easily misidentified. A basic understanding of the potential and common appearances of lysosomes and autophagosomes is critical for TEM analysis. Here, we describe characteristics that should be assessed to properly identify autophagic organelles and provide recommendations for effective classification (Supplementary Figure 1).

Our ultimate goal was to identify and quantify difficult-to-measure autophagic machinery in clear terms and present a novel approach to measure all cellular degradation machinery using free, open-source software. These techniques can be used to reproducibly quantify and characterize changes in the organelles associated with autophagy. Furthermore, for researchers who wish to perform more complex calculations and resource-intensive imaging, we also offer a protocol for 3D reconstruction of recycling machinery.

## 2 Methods and Materials

### 2.1. Mouse Care & Maintenance

Mouse husbandry was performed based on prior protocols [41] according to with protocols approved by the University of Iowa Animal Care and Use Committee (IACUC). Male C57Bl/6J mice were housed at 22 °C with a 12-h light, 12-h dark cycle, and free access to water and standard chow. Mice with a tamoxifen-inducible knockout of DRP1 in skeletal muscle were generated by crossing mice carrying a homozygous floxed allele of DRP1 with mice carrying a tamoxifen-inducible Cre recombinase under control of the myogenin promoter (Jackson Lab) in skeletal muscle as previously described [42,43]. Myotubes were isolated from these mice, using protocols described below.

### 2.2. Fly Strains and Genetics

A mitochondrial assembly regulatory factor (Marf) knockdown fly was generated according to previous protocols [44]. Genetic crosses were performed on yeast corn medium at 22 °C. W1118 flies were used as genetic background controls. Mef2-Gal4 (III) was used to drive muscle-specific Marf RNAi (BS# 55189) to achieve gene knockdown. Mef2-Gal4 (BS# 27390) stocks were obtained from the Vienna Drosophila Stock Center and Bloomington Drosophila Stock Center. All chromosomes and gene symbols are as mentioned in Flybase (http://flybase.org).

### 2.3. Isolation of Satellite Cells & Differentiation

When adopting this protocol, an individual who was blinded to the mouse genotype or treatment conducted the experiment, including isolation, differentiation, and fixation of murine and human cells. This individual did not perform later analyses to mitigate bias. Satellite cell isolation and differentiation for thapsigargin treatment and DRP-1 ablation were performed as described previously, with minor modifications [11,41,45]. When C57B1/J1 mice reached 8–10 weeks of age, mice were anesthetized using isoflurane. Skeletal muscles of the gastrocnemius and quadriceps were excised and washed twice with 1× phosphate-buffered saline (PBS) supplemented with 1% penicillin-streptomycin and 0.3% fungizone (300 μL/100 mL). Dulbecco’s modified Eagle’s medium (DMEM)-F12 media with 0.2% collagenase II (2 mg/mL), 1% penicillin-streptomycin, and 0.3% fungizone (300 μL/100 mL) was added to the muscles and shaken for 90 min at 37 °C. This media was removed, muscle was washed with PBS x4 times, and media replaced with DMEM-F12 media containing 0.05% collagenase II (0.5 mg/mL), 1% penicillin-streptomycin, and 0.3% fungizone (300 μL/100 mL), before shaking for 30 min at 37 °C. Tissue was then ground until all cells were dislodged from the tissue matrix and were passed through a fine, 70-μm cell strainer. Isolated cells were centrifuged, resuspended, and plated on BD Matrigel-coated dishes. Adherent cells were differentiated into myotubes by adding DMEM-F12, 20% fetal bovine serum (FBS), 0.004% (40 ng/mL) basic fibroblast growth factor (R&D Systems, 233-FB/CF), 1× non-essential amino acids, 0.14 mM β-mercaptoethanol, 1× penicillin/streptomycin, and 0.3% fungizone (300 μL/100mL). Myotubes were maintained in medium containing 0.001% (10 ng/mL) growth factor until reaching 85% confluency, then were differentiated in DMEM-F12, 2% FBS, and 1× insulin–transferrin– selenium.

### 2.4. Human Myotubes

GIBCO^®^ Human Skeletal Myoblasts from ThermoFisher Scientific (A1255) were thawed and plated in HG DMEM containing 1% penicillin/streptomycin, 1% fungizone, and 2% horse serum. Cells were differentiated after 48 h, and myotubes were extracted.

Fibroblasts grown *in vitro* to the third passage were plated in 6-well tissue culture plates (5 × 10^5^ cells per well) in DMEM (Invitrogen) supplemented with 10% heat-inactivated fetal bovine serum, 100 U/ml penicillin, 100 μg/ml streptomycin, 0.25 μg/ml fungizone, 1 mm sodium pyruvate, and 10 mm HEPES at 37 °C in a humidified incubator with 10% CO_2_. Cells were infected with Ad-Cre and Ad-GFP was a control.

### 2.5 Immunogold Labeling

Immunogold labeling was performed as previously described [46]. Ultrathin cryosections were prepared, and single- or double-immunogold labeling was performed using antibodies and protein A coupled to gold. After labeling, sections were imaged via TEM. Specifically, primary skeletal myotubes were fixed for 1 h in 4% paraformaldehyde (PFA) and 0.1% glutaraldehyde in 0.1 M phosphate buffer. From there, vibratoming of 40-50 μm section into 0.1M phosphate buffer was performed. Each section was washed 3 times for 10 minutes each with the same 0.1 M phosphate buffer. For the first blocking step, 0.1% NaBH4 in 0.1 M phosphate buffer was applied for 15 minutes. After blocking, phosphate buffer was used for washing 4 times for 10 minutes each time. To permeabilize, 0.05% Triton X-100 in 0.1M phosphate buffer was applied and incubated for 15 mins at 30 °C. After this, phosphate buffer was used for washing 3 times for 10 minutes each time. For the second blocking, Aurion Blocking Solution was added and the solution was incubated at room temperature for one hour. Incubation buffer was used to wash after blocking, again in triplicates of 10 minutes.

From there, 1-5 μg/ml of the primary antibody in incubation buffer was applied and incubation at 4 °C occurred overnight. For mitofusin-1, Anti-Mitofusin 1 antibody was used (Abcam; ab126575); for CAV-1, caveolin-1 antibody was used (Cell Signaling Technology; 3238); and for LC3, LC3B (D11) XP^®^ Rabbit mAb was used (Cell Signaling Technology; 3868). After overnight, the incubation buffer was used to rinse 6 times over, incubating for 10 minutes each time. From there, secondary antibody incubation occurred with 1:100 ultrasmall gold conjugated secondary antibody in incubation buffer overnight at 4 °C. After overnight, again the incubation buffer was used to rinse 6 times over, incubating for 10 minutes each time. Furthermore, 3 washes, each 10 minutes, of PBS were also used. From there, post-fixation occurred through incubation in 2% Glutaraldehyde in 0.1M phosphate buffer for 2 hours. After time elapsed, phosphate buffer was utilized for washing across 4 times, 10 minutes each time. It was quickly rinsed with distilled H_2_O for 15 seconds, three times. For silver enhancement, AURION SE-EM silver enhancement solution was applied, and sample incubated for 45 minutes at room temperature. From there, it was quickly rinsed with distilled H_2_O for 15 seconds, three times. Phosphate buffer was also utilized for washing across 4 times, 10 minutes each time. Once washed, to perform osmification, 0.5% OsO4 in 0.1M phosphate buffer was added for 15 mins. After this, the sample was finally washed in the same phosphate buffer for 10 minutes twice.

From there, dehydration and embedding in resin was performed. To do so, first the sample was again placed in 4% PFA and 0.1% Glutaraldehyde for 1 h. From there, they were washed with phosphate buffer three times for 20 minutes. Following this, a graded ethanol wash was performed at 15 minutes each progressing from 25% to 50% to 75% ethanol. 95% ethanol was finally used for wash for 30 minutes. From there, the mixture was washed and replaced with an 1:1 mixture of 95% ethanol and LR white resin for 1 hr. Finally, a pure 100% LR white resin solution was added. After an hour, it was replaced with a new 100% LR white resin which again incubated at room temperature for 1 hr. This sample was cured under UV overnight with a vacuum to remove any excess liquid. 90 nm ultrathin sections were obtained and imaging occurred as described for TEM samples below.

### 2.6 Lysotracker

The protocol was performed as previously described [47]. LysoTracker^™^ Red DND-99 (ThermoFisher Scientific, L7528) was diluted to a final concentration of 1 mM with dimethyl sulfoxide (DMSO) to create a stock solution, which was then mixed with warm growth media at a 1:2000 dilution. Growth media was aspirated from cells and replaced with the working LysoTracker^™^ Red DND-99 solution. Cells were imaged live using an SP-8 confocal inverted microscope with a visible light laser at a 577 nm excitation wavelength and a 590 nm ± 10 nm emission wavelength, which allowed a yellowish pseudo coloration to be observed. To stain fixed cells, cells were grown in culture media on a #1.5 cover glass, either embedded into a petri dish or divided by plastic-walled growth chambers to optimize microscope optics. Cells were incubated for 30 min with a LysoTracker™ Red DND-99 working solution. The staining solution was then aspirated from the plate, rinsed, and subsequently fixed in 4% PFA. Confocal image stacks were captured with a Zeiss LSM-5, Pascal 5 Axiovert 200 microscope, using LSM 5 version 3.2 image capture and analysis software and a Plan-APOCHROMAT 40x/1.4 Oil DIC objective. Images were deconvoluted with National Institutes of Health (NIH) ImageJ software and BITPLANE-Imaris software. Imaris software analysis was used to measure lysosome number, volume, and area. Experiments were conducted in triplicate, at minimum, and 10–20 cells per condition were quantified.

### 2.7. Immunofluorescence

Immunofluorescence was performed as previously described [41,48]. For, live-cell imaging, live cells were plated and imaged in MatTek 35 mm glass-bottom culture dishes and grown on Matrigel. After growth, the cells were fixed with 4% (w/v) PFA in PBS for 30 min, then permeabilized with 0.25% Triton X-100 in PBS for 10 min at room temperature. Fixed cells were then blocked with 10% bovine serum albumin in PBS and incubated with rabbit anti-lysosomal associated membrane protein (LAMP-1; Cell Signal: D2D11) antibody in 1% BSA in PBST (PBS + 0.1% Tween 20) at a 1:25 dilution at 4 °C overnight. After three PBS washes, each 5 minutes long, Alexa Fluor 488-conjugated goat-rabbit mouse IgG (Life Technologies: A-11008) secondary antibodies were added at 1:1000 dilution in 1% BSA and incubated at room temperature for 45 minutes in the dark. After another three PBS washes, coverslips were mounted onto glass slides with ProLong Diamond Antifade with 4’,6-diamidino-2-phenylindole (DAPI) and allowed to dry overnight. Confocal image stacks were captured with a Zeiss LSM-5, Pascal 5 Axiovert 200 microscope, using LSM 5 version 3.2 image capture and analysis software and a Plan-APOCHROMAT 40x/1.4 Oil DIC objective. Imaris software analysis was used to measure lysosome intensity, length, and sphericity. Experiments were performed in triplicate, at minimum, and 10–20 cells per condition were quantified.

### 2.8. Thapsigargin Treatment

Fibroblasts, human myotubes, and mouse myotubes were treated with thapsigargin (2 μg mL^-1^; Sigma) for 10 h, followed by crosslinking with Trump’s fixative [49] with 4% PFA and 1% glutaraldehyde for 10 min as previously described [11,50].

### 2.9. TEM Processing of Myoblasts, Fibroblasts, and Myotubes

The cell types followed a near identical procedure. Myoblasts and fibroblasts were isolated according to the above methods and placed in six-well poly-D-lysine–coated plates for TEM processing.

Myotubes were cultured on Matrigel coated plates. For 1h, cells were incubated at 37 °C with 2.5% glutaraldehyde in 0.1 M sodium cacodylate buffer. This resulted in cell fixation. From there, after rinsing twice with 0.1 M sodium cacodylate buffer, secondary fixation at room temperature for 30 minutes to one hour occurred using 1% osmium tetroxide and 1.5% potassium ferrocyanide in 0.1 M sodium cacodylate buffer.

After secondary fixation, a five-minute washing with 0.1 M sodium cacodylate buffer (7.3 pH) occurred. From there, two washings of five minutes with diH_2_O ensured the plates were cleaned. While keeping all solutions and plates at room temperature, the plates had 2.5% uranyl acetate, diluted with H_2_O, added and were incubated overnight at 4° C. Following this, dehydration was performed though an ethanol gradient series. After dehyrdation, the ethanol was replaced with Eponate 12_™_ mixed with 100% ethanol in a 1:1 solution. The cells were allowed to incubate, again at room temperature, for 30 minutes. This was repeated three times, for an hour each time using 100% Eponate 12_™_. The plates were finally placed in new media and placed in an oven overnight at 70 °C.

The plates were cracked upon hardening, and the cells were separated by submerging the plate in liquid nitrogen. An 80nm thickness jeweler’s soul was used to cut the block to fit in a Leica UC6 ultramicrotome sample holder. From there, the section was placed on formvar-coated copper grids. These grids were counterstained in 2% uranyl acetate for 2 minutes. Then these grids were counterstained by Reynold’s lead citrate for 2 minutes. Images were acquired by TEM on either a JEOL JEM-1230, operating at 120 kV, or a JEOL 1400, operating at 80kV.

### 2.10. Systematic ImageJ Parameters and Measurement

Using documented parameters and quantification methods [11], a unique individual imaged the entire cell at low magnification. Obtained images were uploaded to ImageJ in an acceptable format, such as TIFF. The cell was then divided into quadrants using the ImageJ quadrant picking plugin (*https://imagej.nih.gov/ij/plugins/quadrant-picking/index.html*, accessed August 21, 2021) to ensure random and unbiased quadrant selection for quantification. After sectioning the image into four quadrants, two quadrants were randomly selected for complete analysis. Three independent, blinded individuals quantified these quadrants as described below. Their collective findings were averaged to decrease individual subjective bias. To ensure accurate and reproducible values, measurements were repeated on a minimum of 10 cells each. In the future, if significant variability is observed among the individuals performing the analysis, increasing the sample number (n) by expanding the number of cells quantified was found to decrease variability.

All analysis methods were developed using NIH ImageJ software. Necessary measures should be set on ImageJ prior to analysis (Analyze > Set Measurements: Area, Mean gray value, Min & Max gray value, Shape descriptors, integrated density, Perimeter, Fit ellipse, Feret’s Diameter). Lysosomes, autolysosomes, LDs, and autophagosomes, were measured, including area, circularity, and length using the Multi-Measure region of interest (ROI) tool in ImageJ based on established measurements [11,51]. Using the freehand tool in NIH ImageJ 1.49, we manually traced the cellular degradation machinery membrane to determine area or volume. A 19 × 23 cm rectangular grid was overlaid on each image to quantify cellular degradation structures and the numbers were presented per 10 μm^2^ of cytoplasm.

### 2.11. Statistical Analysis

Results are presented as the mean ± standard error of the mean. Data were analyzed using unpaired Student’s T-tests. If more than two groups were compared, one-way analysis of variance (ANOVA) was performed, and significance was assessed using Fisher’s protected least significance difference test. For T-tests and ANOVA, the GraphPad and Statplus software packages were used (SAS Institute, Cary, NC). For all statistical analyses, significant differences were accepted when p < 0.05.

### 2.12 3D reconstruction of cellular degradation components using Amira

Following isolation of myotubes in a flex fixative, serial block face-scanning electron micrscopy was performed according to established protocols [16]. Once isolated, *Amira* 3D reconstruction (ThermoFisher Scientific; Waltham, Massachusetts) software was utilized to perform 3D reconstruction of machinery according to the protocol found in Section 3. All videos were created in the *Amira* software program according to Garza-Lopez et al. 2022 [16].

## 3 Protocol

### TEM PROTOCOL

3.1. Downloading and Preparing ImageJ Software for Analysis

3.1.1. Download ImageJ software from the official NIH website (https://imagej.nih.gov/ij/download.html).

3.1.2. Install and open the ImageJ software.

3.1.3. Select Analyze ➧ Tools ➧ ROI Manager to open the ROI Manager, which is used to record and track measurements.

3.1.4. Click on Analyze ➧ Set Measurements to input the measurements for ImageJ to perform, such as area, circularity, and perimeter.

3.1.4.1. For the current protocol, area and count were the focus; however, all available measurements may be used, depending on the study aims.

3.1.5. Import image to be analyzed directly into ImageJ. A TIFF or DM3 file is recommended to provide high-quality.

3.1.5.1. Alternatively, click File ➧ Open to open the selected image.

3.1.6. Considerations

3.1.6.1. For accuracy and reproducibility, ensure that each image contains a scale bar, bar length, and image magnification. The scale bar and bar length are important for setting the appropriate units in the ImageJ settings.

3.1.6.2 Quantification of samples should be performed by three individuals in a randomized and blinded manner to ensure an unbiased approach.

3.1.6.3. To save time, images may be divided into quadrants, and same quadrants should be analyzed across all images.

3.2. Analyzing Lysosomes, Autophagosomes, and Autolysosomes (Supplementary Figure 2A–C)

3.2.1. Click on Freehand Selections to access the Freehand tool.

3.2.2. Trace the outline of the entire cell.

3.2.3. Click Add on the ROI Manager. This ROI will be used to normalize later measurements.

3.2.4. To obtain length and width, use the Straight-Line tool to draw a line down the major and minor axes of each organelle (Supplementary Figure 2A–C, Step 1).

3.2.5. Trace the membrane of each lysosome, autophagosome, or autolysosome. Add the shape to the ROI Manager (Supplementary Figure 2A–C, Step 2).

3.2.6. Click Measure in the ROI Manager to obtain the area measurements.

3.2.7. Add the measurements to the ROI Manager and use the Measure function to obtain numerical values for each measurement.

3.2.8. Considerations

3.2.8.1. Ensure that autophagosomes, lysosomes, and autolysosomes are measured separately because the ROI Manager will group all functions for statistical analysis.

3.2.8.2. The number of autophagosomes, lysosomes, or autolysosomes counted in the cell should be normalized against the total cell area.

3.3. Analyzing Lipid Droplets

3.3.1. Repeat Steps 2.1–2.5 for LDs to obtain basic measurements needed for analysis (Supplementary Figure 2D).

3.3.2. For each cell, calculate total area of all LDs. The amount of lipid coverage is the total area of all LDs divided by total cell area.

3.3.2.1. This process can be used to determine the percent coverage of other subcellular structures, including mitochondria and recycling machinery.

3.3.3. Contact sites between organelles can be measured by first using the Freehand tool to trace the outer membranes of both subcellular structures being analyzed, as described in Step 2.4 (Supplementary Figure 2D, Step 1).

3.3.4. To determine the contact site length, click on the Straight, Segmented, or Freehand Lines tool on the toolbar, and select Freehand Line. Draw a line spanning the length of the contact site, add the measurement to the ROI Manager, and use the Measure function to determine the contact length (Supplementary Figure 2D, Step 2).

3.3.5. Contact distance may be similarly measured using the Freehand Line tool to draw a line between two objects being measured.

3.3.6. Calculate percent coverage by dividing the cumulative contact lengths by the percent coverage of one of the two subcellular features in question, as determined in Step 3.2. Multiply the value by 100 to obtain a percentage.

### 3D RECONSTRUCTION PROTOCOL (UTILIZING WACOM TABLET)

3.A. Download Amira software from https://www.thermofisher.com/us/en/home/industrial/electron-microscopy/electron-microscopy-instruments-workflow-solutions/3d-visualization-analysis-software/amira-life-sciences-biomedical.html (accessed on 4 August 2022) and open.

3.B. Transfer all orthoslices to be analyzed from **Project View > Open Data.** Ensure all are transferred by selecting **Read Complete Volume into Memory**.

3.C. Navigate between the **Project** subsection, to select images to analyze and the the **Segmentation** subsection, to alter segmentation tools. We recommend the **Brush** tool with a size of 2.

3.D. Calibrate the Wacom Pen and open Amira on the Wacom tablet in accordance with prior protocols [16].

3.E. Using arrow keys, scroll through orthoslices until desired recycling machinery is found. It is recommended to verify its identity by looking at additional orthoslices. For each organelle, outline with a material and press F to segment area. Alter material for each organelle to pseudo color organelles.

3.F. Repeat this process for each ortho slice that organelle appears on, making sure that all independent organelles consistently use the same material color. Later on, aspects of these materials, such as color and visibility, can be altered on the **Materials** menu.

3.G. Once all are segmented, on the **Project Menu**, click on **Selection Labels > Generate Surface** and select **Apply**.

3.H. In the workplace area of Amira, rename the newly generated box with the “**. surf”** suffix and click **Surface View**. Toggle orthoslice and 3D reconstruction materials on and off to switch between isolated and overlay view.

3.I. Make scale bars by right-clicking in the gray area under the **Project** subsection and selecting the option **Scalebars**. Adjust scale bars as necessary, with a readable width and font, as well as only leaving x-axis, unless y-axis scale is also necessary.

3.J. Previously [16], mitochondrial quantifications performed has included volume, 3D area, and length. Furthermore, mitochondrial complexity index is measured by SA^3^/16pi^2^V^2^ while mitochondrial branching index measures transverse branching divided by longitudinal branching [52]. While created for mitochondria, these same quantifications can be used as parameters for recycling machinery. Additionally, recycling machinery has other quantifications that may be applied including mitochondrial-lysosome or lipid droplet-mitochondrial interactions distances and surface area interactions.

## 4 Result

This protocol describes a method to obtain reproducible measurements and identify structures involved in autophagy. Below, we show the results obtained using this TEM image analysis approach.

### 4.1. Identification of organelle compartments by immunogold labeling

With the pitfalls associated with correctly identifying organelles by TEM morphology alone, other methods may be required to confirm organelle identity. One of the most effective alternatives is immunogold labeling used in electron microscopy to analyze organelle marker proteins. As a positive-control, it is useful to perform immunogold labeling on a easily identifiable organelle. For example, mitochondrial GTPase proteins, mitofusin 1 and 2 (MFN1 and MFN2), function in mitochondrial fusion reactions [35,36,53–56] and MFN1 is used to identify mitochondria in tissues as MFN1-positive puncta (Figure 2P–Q).

Immunogold labeling can also be used to identify organelles associated with autophagy. Many novel yeast genes that are essential for autophagy (autophagy-related, or ATG genes) have been characterized, and most of their mammalian homologs have been identified [57]. Microtubule-associated protein 1 light chain 3 (LC3), the mammalian homolog of Atg8 [58], is a reliable marker for mammalian autophagosomes, which can be identified by the formation of LC3 labeling along the limited membrane (Figure 2R–S). LC3 expression in autophagosomes (Figure 1, blue arrows) and phagophores (not shown) may vary due to LC3 degradation by lysosomal hydrolases, making it a challenge to identify late-stage autophagic materials [59]. Moreso, since autophagosomes are three-dimensional organelles and are being imaged in two-dimensions. However, identifying LC3-positive puncta is still valuable to identify autophagosomes. Immunogold labeling has also been performed with caveolin-1 (CAV-1), a marker protein for specialized membrane domains known as caveolae, which ultimately accumulate in caveosomes that mature into MVBs upon endocytosis [60]. Therefore, CAV-1 immunogold labeling can be used to identify vesicles (Figure 2T–U) and the presence of CAV-1 puncta in an ROI excludes those vesicles from classification as autophagosomes, indicating instead a vesicle such as, multivesicular or multi-inclusion body. After testing immunogold labeling, we examined changes in cellular degradation machinery under other conditions.

### 4.2. Thapsigargin treatment alters lysosome, autolysosome, and autophagosome morphology

Thapsigargin is a sarcoplasmic-ER Ca^2+^-ATPase (SERCA) inhibitor, that decreases the length of mitochondria–ER contacts in treated cells, while also inducing ER stress [11,61]. We investigated morphological changes in lysosomes, autolysosomes, and autophagosomes in response to thapsigargin (Figure 3) using our TEM image analysis protocol. We found that mean lysosomal area and the number of lysosomes per square micron significantly increased in response to thapsigargin treatment in primary mouse skeletal myotubes (Figure 3E–F). The mean area of autolysosomes and the number of autolysosomes per square micron had an even greater increase than seen in lysosomes (Figure 3G–H). The mean autophagosomal area and number of autophagosomes per square micron also significantly increased in thapsigargin-treated cells (Figure 3I–J). Similar results were seen in mouse fibroblasts (Figure 3K–T) and human myotubes (Figure 3U–AD). Human myotubes displayed the largest increases in autophagy recycling machinery of all assessed components. These quantifications are shown with representative images for each cell type (Figure 3A–D, K–N, and U– X). The ability of thapsigargin to inhibit ER function and promote cell stress support a model in which cell-stress-induced organellar damage increases lysosome and autophagosome degradation of damaged organelles. Thus, morphological changes detected and quantified using our TEM method are consistent with the expected effects of thapsigargin treatment.

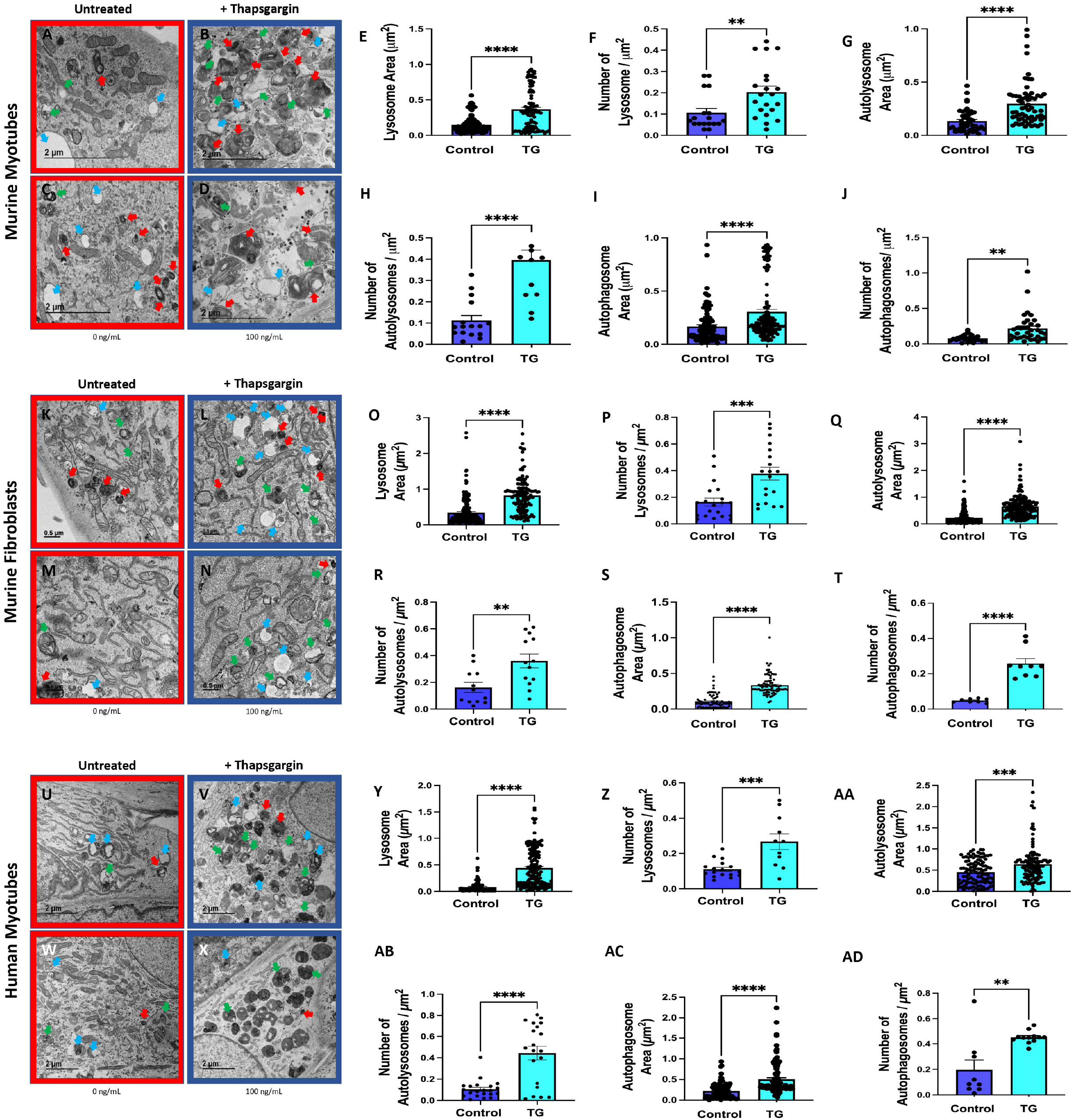

### 4.3. DRP-1 ablation results in increased degradation machinery

Dynamin-related protein (DRP-1) is a crucial regulator of mitochondrial fission [43,62]. Previous studies found that impaired mitochondrial fission can have downstream effects on organellar morphology and function throughout the cell [63]. In the absence of DRP1, mitochondria undergo fission less frequently, resulting in longer mitochondria that can trigger downstream effects, including apoptosis [62]. To test our method, we generated a skeletal muscle-specific *Drp1* knockout mouse and noted changes in the degradation machinery. Specifically, our study focused on lysosomes, autophagosomes, and LDs, which are all closely linked to the autophagy process. *Drp1* ablation in a skeletal myotube-specific knockout model (DRP-1smKO), resulted in significantly more lysosomes than in wild-type controls (Figure 4A–F, red arrows). We also found increased lysosome numbers per square micron and lysosomal area per square micron, although the change in lysosomal area was not as great as the change in lysosome number (Figure 4G–H). Similarly, DRP-1smKO also showed a significant and large increase in autophagosome number over wild-type controls (Figure 4I–L, red arrows). We also observed increased autophagosome number per square micron and autophagosome area per square micron, although the change in area was less than the change in number (Figure 4M–N). Based on changes in percentage and degree of significance, the autophagosome increase was greater than the lysosome increase, suggesting that reduced mitochondrial fission may cause larger shifts in cargo vessel formation than in lysosome formation, although both organelle types increased significantly. Autophagosome-lysosome fusion events may also contribute to this disparity, as intermediate fusion phase structures more closely resemble autophagosomes than lysosomes.

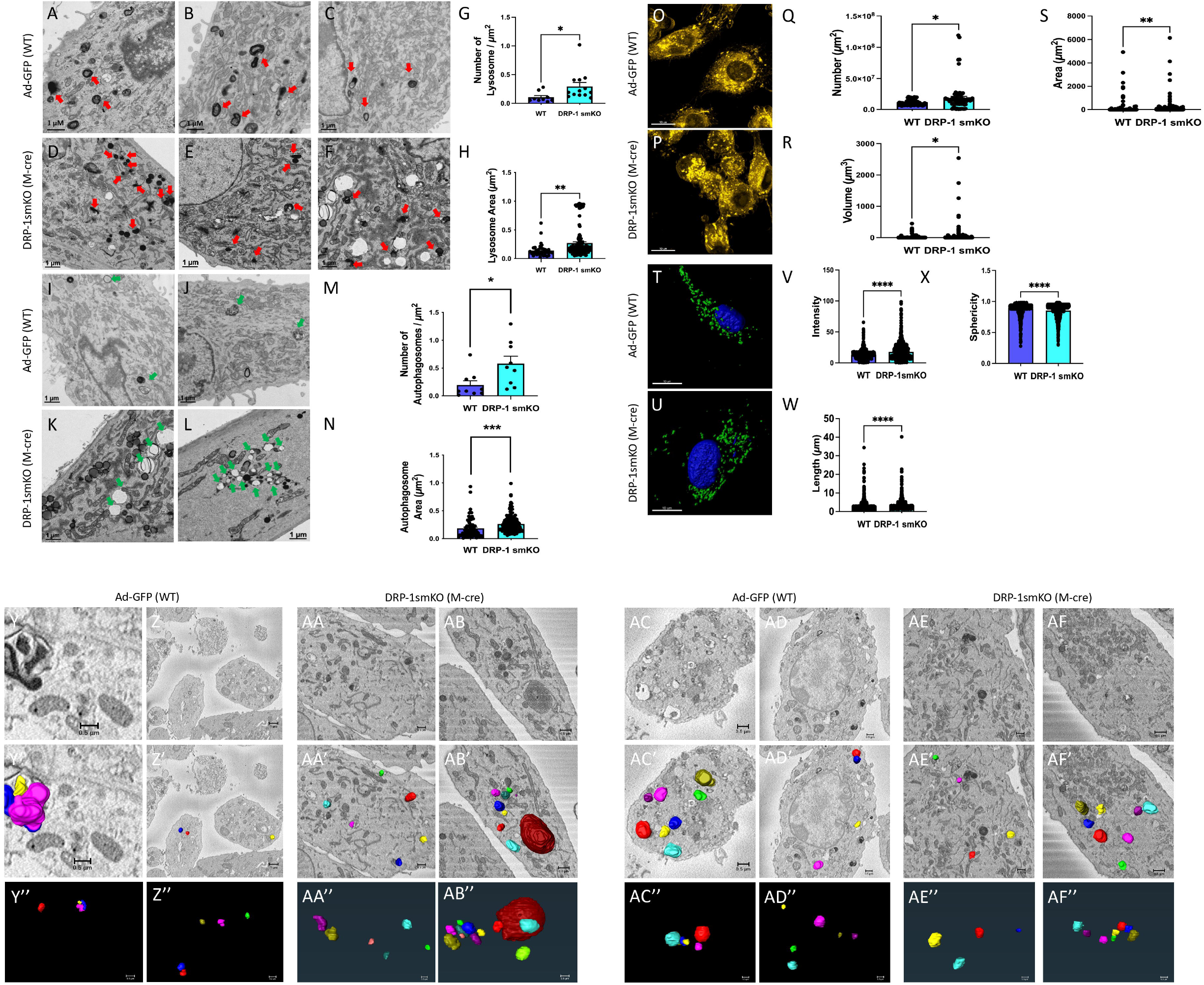

To validate these results, we used the fluorescent dye, LysoTracker, to image lysosomes in DRP-1smKO myotubes because its ability to label and track acidic organelles in live cells allows it to effectively identify highly acidic lysosomes. Similar to TEM data, the LysoTracker assay showed significantly more lysosomes in DRP-1smKO myotubes than in wild-type controls (Figure 4O-P). Increased lysosome number, calculated lysosome volume, and lysosomal area were also observed in DRP-1 knockout mice (Figure 4Q–S). These key quantitations are similar to those determined by TEM analysis; however, LysoTracker provided better certainty of lysosome identity and allowed use of traditional statistical analyses to determine lysosome area and numbers. Lysosomes can also be identified by LAMP1 immunostaining (Figure 4T–U) [64], which correlates with the number and size of active lysosomes (Figure 4V). While relative intensity can estimate LAMP1 expression, lysosome area measurements are not reliable by LAMP staining. Other metrics, including length and sphericity, can be determined using these fluorescent dyes, suggesting that lysosomal dysfunction occurs as length increases and sphericity decreases (Figure 4W–X).

To further validate these results and see if the 3D structures of recycling machinery showed significant differences upon loss of DRP-1, we also provide representative 3D reconstruction of both lysosomes (Figure 4Y-AB”; SV1-4) and autophagosomes (Figure 4AC-AF’’; SV 5-8). The top image shows a representative orthoslice of the region of interest (Figure 4Y-AF). Once segmentation was performed, organelle 3D reconstructions were overlaid on the orthoslice (Figure 4Y’-AF’). To allow for better visualization, we also show isolated 3D reconstruction on the X-by Z-plane to allow for their depth to better be visualized (Figure 4Y’’-AF’’). However, static images of 3D reconstruction may not capture the scope of 3D reconstruction; therefore, we also provide associated videos showing organelles for all representative images (SV 1-8). Our study shows that lysosomes appear larger in DRP-1 smKO (Figure 4AB’’). Additionally, upon loss of DRP-1 lysosomes appear to be closer together in 3-dimensional spatial orientation (Figure 4AA”-AB”). Autophagosomes appear to have a less significant alteration, but they do appear to be more elongated in the z-axis upon DRP-1 smKO (Figure 4AC”-AF”), a detail that would otherwise not be observed in TEM.

### 4.4. DRP-1 ablation results in increased lipid droplets

We also measured LDs in skeletal muscle from DRP-1smKO mice, which had significantly more LDs than WT controls (Figure 5A–B, red arrows). We observed a large increase in lipid area and number of LDs per square micron (Figure 5C–D). Based on percent change, the LD increase was larger than the observed increases in both lysosomes and autophagosomes following DRP-1 ablation (Figures 4 and 5). To see if spatial relationships of LDs changed upon DRP-1 loss, we also looked at SBF-SEM 3D reconstruction of LDs. For both WT and DRP-1smKO mice we show a representative x-by y-plane orthoslice (Figure 5E-H), 3D reconstruction overlaid on orthoslice (Figure 5E’-H’), isolated 3D reconstruction showed on a slightly different plane, the X-by Z-plane, to allow for depth to better be visualized (Figure 5E”-H”), and video of isolated 3D reconstruction (SV 9-12). We observed that after DRP-1 loss, LDs were much more clumped, both on the x-and y-axis, but also on the z-axis, and lipid droplets appeared to be larger sizes (Figure 5G”-H”).

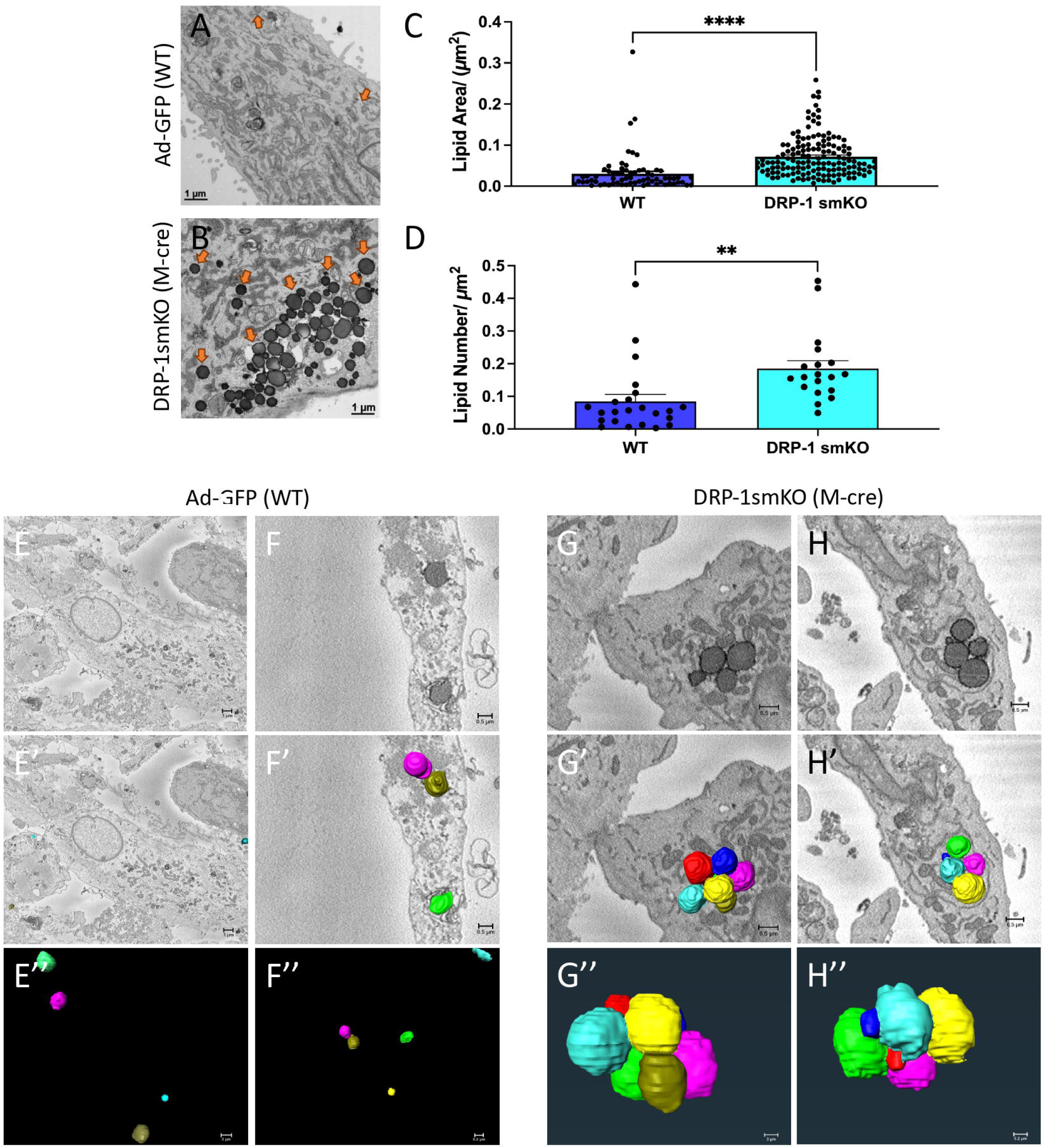

### 4.5. Knockdown of Marf resulted in more abundant lysosomes

In addition to DRP-1, we sought the quantify lysosomal changes in response to knockdown of other key mitochondrial proteins. Mfn2 is an important regulator of mitochondrial fusion [65,66] and Mfn2 deficiency has been associated with disrupted ER morphology and mitochondria–ER contacts, resulting in dysfunctional calcium signaling [65,66]. Loss of Mfn2 was also recently shown to influence autophagic pathways [65–68] by stalling autophagy at the lysosome and autophagosome stages, causing a buildup of both autophagosomes and lysosomes by inhibiting their fusion [68]. The *Drosophila* homolog of Mfn2 is Marf, and knockout of genes upstream of Marf had downstream effects on autophagy [67]. Given this emerging link between autophagy and Marf/Mfn2, we examined the effects of Marf knockdown in *Drosophila* tissue, which produced a significant and large increase in lysosome number compared with WT controls (Figure 6A–B, D). We also observed an increase in average lysosomal area (Figure 6C). These findings indicate a potential upregulation of autophagy, which may represent an autophagic response to ER and mitochondrial stress caused by loss of Marf [65–68]. Further research into changes in other cellular degradation machinery following loss of Mfn2/Marf could better elucidate the effects of Mfn2/Marf on autophagy.

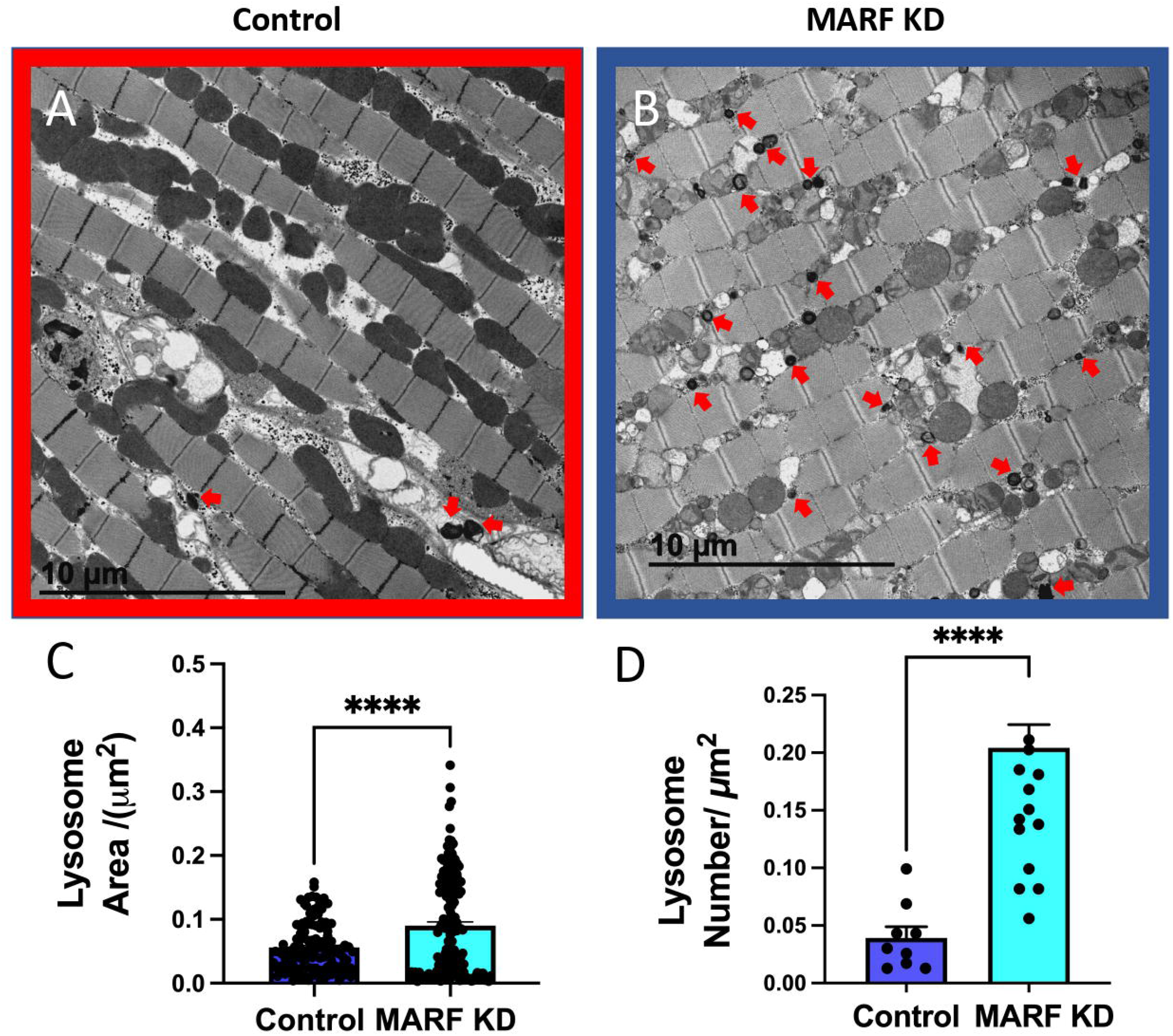

## 5 Discussion

The method we have described involves measuring organelles in each image or image quadrant by defining the area of interest using digital tools, rather than the often-used method of point counting. Point counting can be used to determine the cellular area by laying a grid over the cellular area and determining the distance between gridlines and the number of grid intersections, or points, within the cell. Cellular area may then be estimated using the equation P × d^2^, where p is the number of points and d is the grid distance [40]. Smaller grids can be used to repeat the process for estimating organelle area, and these two values can then be used to determine the percent coverage of organelles. Past studies have used point counting to successfully streamline the process of calculating organelle coverage; however, this method only provides estimations [40]. Even with smaller grid distances, which can increase the calculation accuracy, this approach still requires estimation. The method we described uses ImageJ to more precisely calculate the structure areas. Although using ImageJ is more time-consuming, the results are highly reproducible, generating high-quality data that may be further analyzed using ImageJ. Both analyses require human evaluation and proper identification of the recycling machinery. Both point counting and ImageJ-based measurements are viable ways to measure the frequency of recycling or other types of organelles; however, we believe the accuracy of measurements with ImageJ analysis is worth the increased time commitment [23,24].

An important consideration in the use of TEM is the magnification and scope of cellular degradation machinery components to be considered in these analyses. Significant size heterogeneity is seen among components of the degradation machinery, even within the same classification group, which can vary according to the amount of cargo they carry (Figure 1). For TEM imaging, various organelle types and sizes may require different magnifications (Supplemental Figure 1). Additionally, the purpose of the analysis must be considered when deciding which types of recycling machinery to evaluate, and appropriate statistical analyses should be used (Supplemental Figure 1). For example, the total number of LDs may not be as important as their total cell coverage due to varying LD sizes and proper magnification should be determined based on the necessary measurements to be performed (Supplemental Figure 1). A limitation of our method is that for key measurements, such as area, the total organelle must be visualized to obtain accurate results. Point counting can be used to evaluate images in which the entire organelle is not visible because it relies on estimation [40]. When using ImageJ, the entire organelle is outlined, and magnification that is too high may limit the amount of data that can be collected. However, images of a single cell at varying magnifications may be used by normalizing to a consistent scale across all images.

Although this protocol focuses primarily on evaluation of the degradation machinery, these organelles must not be viewed in a vacuum. Autophagy can target any cell, and many factors can alter autophagy processes in different cells, as seen in cancer, metabolic diseases, or neurodegenerative diseases [1,4,12]. Many organelles are closely associated with the overall autophagy process. Recent research found that omegasomes and autophagosomes primarily form at mitochondria–ER contact sites [39], which may be because the ER phospholipid, phosphatidylinositol 3-phosphate, is needed to activate and form autophagosomes [1–3]. Mitochondrial-derived vesicles can also influence autophagic pathways by transporting proteins and lipids associated with the mitochondria to MVBs [69]. This previously unknown pathway indicates that mitochondria that are not sufficiently damaged to trigger mitophagy can still produce endocytic bundles that are transported to the MVB for recycling via the autophagic pathway. Because cellular degradation machinery can have important effects on organelles, and the inverse is also true, a holistic view is needed to understand the nuances that influence autophagy.

The protocol described here is optimized for statistical analyses; however, it is important to ensure that correct subcellular features of lysosomes, autophagosomes, and LDs are identified and measured. Although organelles can be accurately identified by TEM alone, methods, such as immunofluorescent staining, are recommended in tandem with TEM to achieve clear results, particularly when analyzing lysosomes, autophagosomes, and autolysosomes, which are easily misidentified (Figures 1–2 and 4). Because examining organellar morphology using TEM alone may lead to inaccurate conclusions, we suggest coupling TEM with methods such as immuno-TEM with gold labeling, LysoTracker with correlative light and electron microscopy, and immunohistochemistry or immunofluorescence (Figure 2 and 4) to verify the identity of each structure [5,57,70].

Given the acidity and multitude of proteins associated with lysosomes, there are various ways to identify lysosomes using immunogold labeling, LysoTracker to identify acidic organelles, immunofluorescent dyes to label lysosome-associated proteins such as LAMP1(Figure 4T–U) [64], and indirect immunofluorescence using secondary antibodies bound to a lysosome-associated primary antibody [71,72]. When combining confocal fluorescence imaging and TEM, fluorescence can be used to identify specific proteins and confirm the identities of autophagic organelles (Figure 4O–X), and TEM can be used to measure finer details, including area, average number, and percent coverage (Figure 4A–N). SBF-SEM can from there allow for 3D visualizations and to see details that may not be seen in the 2D plane (Figure 4Y”-AF”; Figure 5E”-H”; SV 1-12). 3D reconstruction can importantly allow for visualization of how organelles exist in relation to each other in a 3D spatial orientation as well as if there is alterations in specifically transverse or longitudinal organelle volumes [16].

Current options to identify and classify autophagosomes are limited. Immunogold labeling can be used to detect LC3, which is currently a commonly used autophagosomal marker that proved effective for our research [57,70,73]. LC3 puncta are not always detectable in autophagosomes; however, immunogold labeling can be used to identify organelles to be excluded from autophagosomes analysis. For example, CAV-1 staining (Figure 2T–U), which is associated with caveolae typically found in MVBs, can identify MVBs that might be mistaken for autophagosomes (Figure 2) [60]. Similarly, perilipin 2, a commonly expressed protein associated principally with LDs, can be used to identify LDs [74]. Future studies that explore new improved immunogold or immunofluorescence labeling options for autophagosomes will be important. Due to the potential ambiguity associated with identifying the cellular degradation machinery, we recommend using at least one additional complementary technique to verify lysosome and autophagosome identity when measuring TEM images. Future studies may also perform quantifications of 3D reconstruction to determine more detailed changes beyond those only shown by TEM.

Using the method outlined here, we have quantified the changes associated with macro autophagy upon treatment with thapsigargin, loss of DRP-1, or knockdown of MARF. Thapsigargin treatment, this is known to cause ER stress and past research has shown that such ER stress promotes lipotoxicity and the formation of lipid droplets [75,76]. Given these previous findings, we believed that autophagy may increase upon thapsigargin treatment in addition to lipid droplets, as previously established for lipid droplets. Specifically, this hypothesized this would come across given that past literature has found that macro autophagy can serve as a protective mechanism for ER stress [77,78]. Our results support the autophagic response to ER stress, as autolysosome, autophagosome, and lysosomes all increased in quantity in three different models (Figure 3). Critically, we also examined size of these organelles, as past literature has also suggested that the size of these organelles relates to their efficiency and function [37,79,80]. In combination, our results show in all three models a large increase in autophagy machinery. ER stress is typically caused by dysregulation of protein folding, causing dysregulation of ER homeostasis [77]. In response, the unfolded protein response is activated, which can have a downstream effect of increasing autophagy to remove defective organelle and macromolecules [81]. Similarly, loss of DRP-1 also was demonstrated to cause an increase in autophagy as seen by increases in size and quantity of organelles and increased LAMP1 expression (Figure 4), and autophagy likely increases through a similar mechanism. Increased LDs have previously been described as a downstream effect of autophagy, consistent with the conclusion that autophagy occurs more frequently following DRP-1 ablation [32]. Increased autophagy may be due to dysfunctional regulation of mitochondrial length, which is seen in response to loss of DRP-1 regulated fission [32,62]. These results suggest that DRP-1 ablation and the resulting lack of mitochondrial fission increase autophagy in cells, demonstrated by upregulation of the cellular degradation machinery. In past studies, mice lacking DRP-1 have had increased accumulation of damaged mitochondria [63], which may induce increased mitophagy and relevant recycling machinery. While DRP-1 is typically associated for regulation of mitochondrial fission [43], past literature has also implicated the loss of DRP-1 with ER stress [82]. Specifically, this may come as a result of loss of calcium homeostasis between the dysfunctional mitochondria and ER. Therefore, this is a possible reason for why loss of DRP-1 mimics the changes seen upon thapsigargin treatment. Similarly, past research has also shown that MFN-2 increases ER stress [83]. found that loss of MARF, the fly homolog of MFN2, increased autophagic organelles, further showing a potential increase in autophagy due to ER stress due to the unfolded protein response. Future research may consider studying specificically ER-phagy with this method to better understand how ER-stress activates autophagy, and verify that DRP-1 and MARF loss increase autophagy as a downstream effect of ER stress.

Although limitations of this TEM analysis and SBF-SEM visualization method exist, when combined with other techniques, reliable identification and quantitation of cellular degradation machinery components may be possible. On a broader scale, this method using ImageJ and/or Amira may be applied to other fields with a focus on organelle structure. For example, mitochondria play key roles in many complex diseases, including type II diabetes, cardiomyopathy, and Alzheimer’s disease [35,36,53–56] and autophagy may contribute to these diseases, given its role in mitophagy to clear dysfunctional mitochondria. Use of TEM and ImageJ to study other organelles in conjunction with the precise methodology outlined here to study key autophagic organelles will improve our understanding of the physiology associated with key organelles and their contributions to disease.

## 6 Perspective on Staining

Lysosome stages may look different when using different EM staining procedures, depending on the material used for preparation (e.g., osmium tetroxide) and the type and amount of additives used (e.g., uranyl acetate, lead citrate, and ruthenium red). Depending on the stain used, lysosomal membrane contrast may be altered, affecting the appearance of lysosome-related structures. All EM images shown here used a grid-based staining technique in all procedures (Table 2). The general TEM sample preparation protocol used glutaraldehyde and 1% osmium tetroxide as fixatives [84– 87]. Post staining on TEM ultrathin sections used 5% uranyl acetate for 6 min to increase membrane contrast and Reynold’s Lead Citrate for 3 min to improve resolution of cellular structures [88–90]. Other stains may be used to optimize the experimental purpose, and staining time should be adjusted according to the sample type. Other viable alternatives exist; for example, ruthenium tetroxide is particularly useful when preparing kidney, liver, and prostate tissue [91,92]. Ammoniated ruthenium oxychloride, commonly referred to as ruthenium red, is frequently used as a polycationic dye to stain negatively charged molecules, including polysaccharides, in tissue sections [93–95]. Although ruthenium red is commonly used for fungal staining, when used with osmium tetroxide, a chemical reaction occurs that may increase the contrast of TEM micrographs [93–95]. Regardless of the stain used, the foremost concern should be maintaining consistency in tissue staining. Different staining in the same organism for example, staining separately for lysosomes, should be avoided. Ideally, the same staining solution should be used for all samples, even at different stages, making it possible to compare different times or stages. If staining protocols different from those described here are used, lysosome appearance may vary.

## 8 Author Contributions

Conceptualization, E.G.-L., Z.V., P.K., J.S. (Jianqiang Shao), R.O.P., E.D.A. and A.H.J.; Methodology, E.G.-L., Z.V., P.K., L.V., J.S. (Jianqiang Shao), S.A., M.M., M.B., S.A.M., J.L., T.A.C., J.L.S., R.O.P., E.D.A. and A.H.J.; software, E.G.-L., Z.V., P.K., L.V., T.A.C., J.L.S., R.O.P., E.D.A. and A.H.J.;

Validation, E.G.-L., Z.V., P.K., K.N., L.V., T.A.C., J.L.S., J.S. (Jianqiang Shao), R.O.P., E.D.A. and A.H.J.;

Formal analysis, E.G.-L., Z.V., P.K., K.N., L.V., J.S. (Jianqiang Shao), T.A.C., J.L.S., R.O.P., E.D.A. and A.H.J.;

Investigation, E.G.-L., Z.V., P.K., L.V., J.S. (Jianqiang Shao), H.K.B., A.G.M., T.A.R., T.A.C., J.L.S., B.G., J.S. (Jennifer Streeter), R.O.P., E.D.A. and A.H.J.;

Resources, E.G.-L., Z.V., P.K., S.A.M., T.A.C., J.L.S., R.O.P., E.D.A. and A.H.J.;

Data curation, E.G.-L., Z.V., P.K., J.S. (Jennifer Streeter), H.K.B., A.G.M., T.A.R., R.O.P., E.D.A. and A.H.J.;

Writing—original draft preparation, E.G.-L., Z.V., P.K., K.N., H.K.B., A.G.M., T.A.R., M.B., S.A.M., B.G., T.A.C., J.L.S., R.O.P., E.D.A. and A.H.J.;

Writing—review and editing, E.G.-L., Z.V., P.K., K.N., H.K.B., A.G.M., T.A.R., S.A.M., B.G., T.A.C., J.L.S., R.O.P., E.D.A. and A.H.J.;

Visualization, E.G.-L., Z.V., P.K., K.N., B.G., T.A.C., J.L.S., R.O.P., E.D.A. and A.H.J.;

Supervision, E.G.-L., Z.V., P.K., B.G., T.A.C., J.L.S., R.O.P., E.D.A. and A.H.J.;

Project administration, E.G.-L., Z.V., P.K., T.A.C., J.L.S., R.O.P., E.D.A. and A.H.J.; Funding acquisition, E.D.A. and A.H.J.

All authors have read and agreed to the published version of the manuscript.

## 9 Funding

This work was supported by NIH grants, R01HL108379 and R01DK092065 to E.D.A. UNCF/BMS EE United Negro College Fund/Bristol–Myers Squibb EE Just Postgraduate Fellowship in the Life Sciences Fellowship to H.K.B., the United Negro College Fund/Bristol–Myers Squibb EE Just Faculty Fund, Burroughs Wellcome Fund Career Awards at the Scientific Interface Award, Burroughs Wellcome Fund Ad-hoc Award, the National Institutes of Health Small Research Pilot Subaward 5R25HL106365-12 from the National Institutes of Health PRIDE Program, DK020593, the Vanderbilt Diabetes and Research Training Center for DRTC Alzheimer’s Disease Pilot & Feasibility Program to A.H.J. The NSF grant MCB 2011577I and NIH T32 5T32GM133353 to S.A.M.Acknowledgments

We would also like to thank our undergraduate colleagues Benjamin Kirk and Innes Hicsasmaz for helping to optimize the analysis technique. y

