## Supplemental Figures & Legend for "Systematic Transmission Electron Microscopy-Based Identification and 3D Reconstruction of Cellular Degradation Machinery"

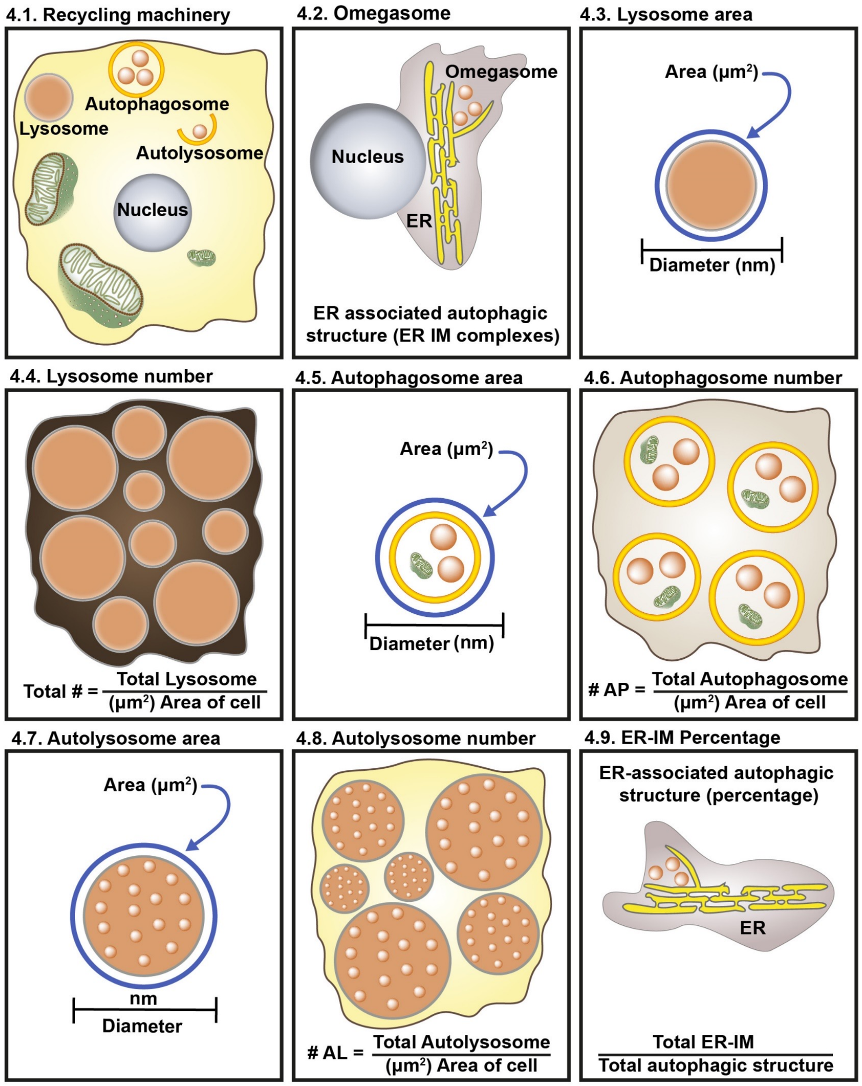


**Supplemental Figure 1. Proper measurement and magnification for analyzing essential recycling machinery.** To accurately assess lysosomes or autophagosomes, the lysosome and autophagosome numbers, area, and diameters should be measured. This strategy is the same strategy applied to mitochondria. (**A–I**) Magnifications should be standardized to acquire a clear understanding of the morphological changes observed in mitochondria and other organelles. Recycling machinery varies considerably in size depending on the organelle. Thus, no single magnification can encompass all structures of interest. For example, omegasomes can be large and require each structure to be individually quantified relative to the endoplasmic reticulum (ER). (**A and B**) The structure and quantity of these organelles vary between cell types and across treatment conditions. (**C–H**) Alternatively, the recycling machinery, such as lysosomes, autophagosomes, and autolysosomes, can be analyzed at 1000× to measure the area, diameter (length and width), and quantity. (**I**) 1000× magnification can be used to count ER-isolation membranes (ER-IMs). To view detailed morphology, the use of magnifications lower than 4000× is recommended.

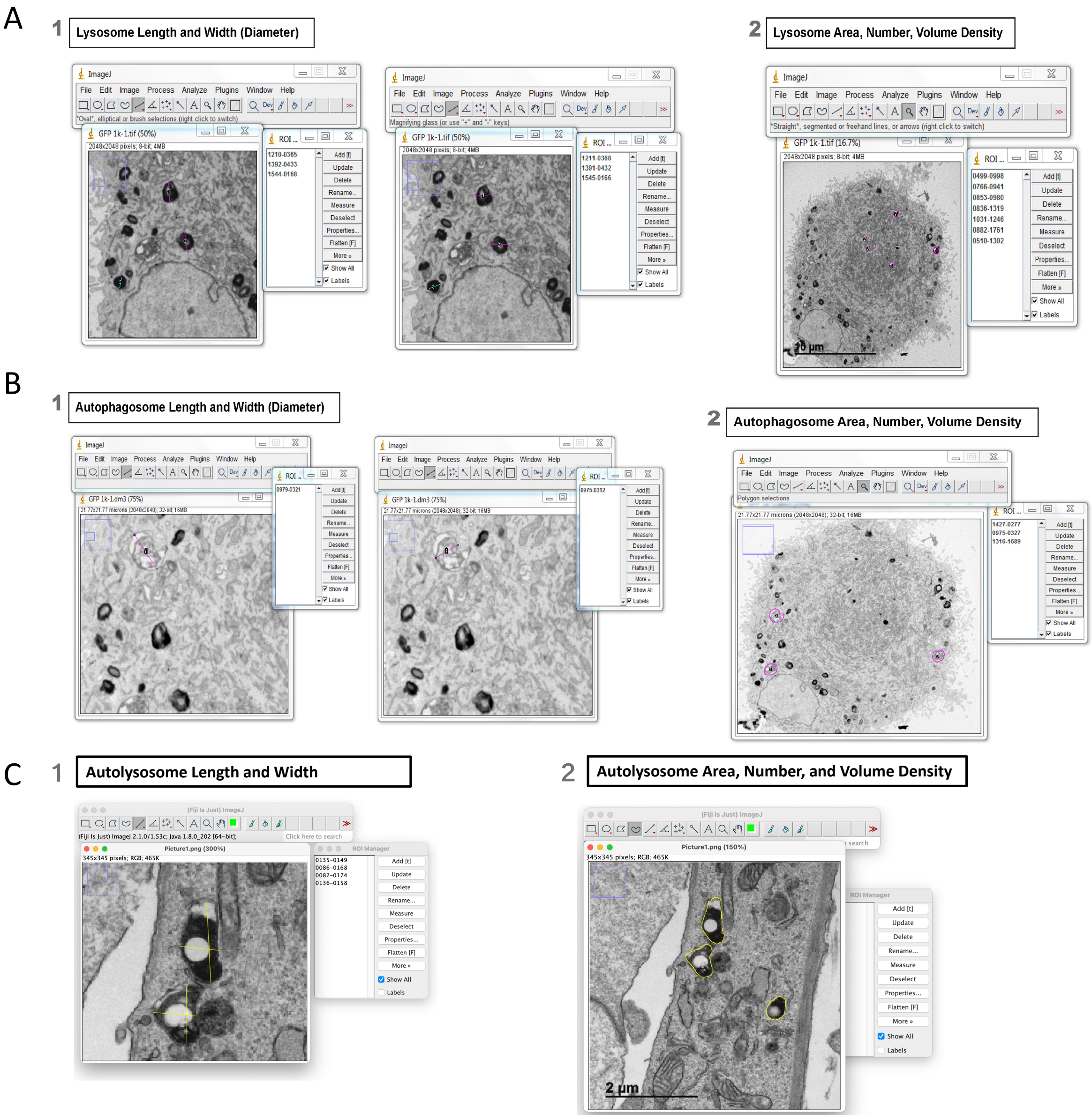


**Supplemental Figure 2. Workflow and representative images of the quantification of autophagosomes, lysosomes, autolysosomes, and lipid droplets.** (**A**) The fundamental process used to evaluate lysosomes using key measurements, including length, width, area, number, and density. Lysosomes are all indicated in representative images by circles quantifying area and lines spanning the Feret's diameters. (**B**) Additional representative images show the same information for autophagosomes and (**C**) autolysosomes. (**D**) Slightly differing quantifications should be performed for lipid droplets, although they may all be easily performed using ImageJ. All of these quantifications should be performed separately, on an organelle type-by-organelle type basis, to avoid the accidental confusion of regions of interest.

It is important to properly identify and distinguish degradation machinery depicted in Figures 1-5. Although under TEM without any other methods being used in tandem, degradation machinery may look similar, several key differences can be used to distinguish these structures.

- Lysosomes typically present as dark objects ranging in size from 0.05 to 1 micrometer, with a median of 0.2 to 0.5 micrometers. Lysosomes and autolysosomes can be misidentified because both share similar circular or elongated, ovular shapes. Additionally, when imaged via transmission electron microscopy (TEM), the membrane will appear as a black circle surrounding both lysosomes and autolysosomes. However, typically primary lysosomes feature a more homogenous appearance that is electron dense throughout. Additionally, autolysosomes typically show evidence of fusion by way of limiting membranes being combined. Lysosomes may feature multiple membranes, which can result in an appearance similar to that of a multilamellar vesicle; however, typically, lysosomes feature fewer membranes and may be smaller, since multilamellar bodies can present as large as 2.5 micrometers. Similarly, MVBs, or late endosomes, can closely resemble lysosomes. MVBs will typically contain content, presenting as small internal vesicles which will also have membranes. Autophagosomes typically present as objects with two visible limiting membranes that range in size from 0.5 to 1.5 micrometers. A key characteristic of autophagosomes that aids in their identification is cytoplasmic cargo they have collected. This can include objects such as clusters of ribosomes or portions of rough ER or mitochondria. The varying stages and cargoes of autophagosomes give them varied appearances, including some with incomplete membranes and "empty" content, whereas others have more complete membranes and contain cargo. At high magnification, autophagosomes appear to have two or more membranes, and the cargo typically appears as defined sacs that are more circular rather than as cristae or other artifacts. This allows for them to be differentiated from autolysosomes, which have a single limiting membrane and limited cargo, or cargo that is already being degraded. At a lower magnification, MVBs may also appear to feature multiple membranes; however, many MVBs contain lipids, which gives the interior compartment of multivesicular bodies an inhomogeneous appearance compared with autophagosomes. Similar to autophagosomes, multivesicular bodies can have a similar cargo-laden appearance, especially from at lower magnification. The cargo in MVBs, however, is typically in more clearly defined vesicles. In such cases, a higher magnification can be used to identify differences between multivesicular bodies and autophagosomes.
- Autolysosomes are typically 0.5 to 1 micrometer and identified by evidence of fusion. They present with a singular membrane and degrading cargo. Multilamellar vesicles can present akin to autolysosomes. Multilamellar vesicles have many lipid bilayers, which give them a ring-like appearance. However, these rings, when examined under high magnification, have a disorganized and crooked pattern that allows for their identification. In contrast, autolysosomes typically have a larger, singular limiting membrane. Additionally, while they have degrading cargo, they also typically present clear, circular, white autophagosomal compartments.
- Lipid droplets are typically homogeneous organelles that can vary greatly in size from 0.01 micrometers to over 10 micrometers, typically around 0.2 to 1 micrometer. Lipid droplets have a very homogenous gray appearance free of any cargo with a weak limiting membrane. However, in some cases, autolysosomes or lysosome can be mistaken for lipid droplets. If lipid droplets present a darker membrane, they can resemble a lysosome. Lipid droplets, however, will have greater diversity in size, a thinner membrane, and more clumping. Autolysosomes can present a similar clear circle in the middle as lipids sometimes do, however lipids do not display the fusion event that autolysosomes do. Lipid droplets are typically located in a single area near the edge of the cell, which is not typical behavior for autolysosomes, which are found closer to the center of cells and rarely clump.

**Supplemental Table 1. Suggested Notes to Consider for the Identification of Degradation Machinery**

### Grid staining for TEM

**Materials**

- 5% Uranyl Acetate (UA)
- Reynold's Lead Citrate
- NaOH pellets
- 1 N NaOH
- 2 clean, disposable Petri dishes
- dd $H_2O$ in squirt bottle
- 3 beakers filled to ¾ full with dd $H_2O$
- Filter paper cut into small triangles
- Self-closing fine tipped forceps
- 2 micro-centrifuge tubes

**Prepare stains**

- Draw up ~ 0.5 ml of clear UA into a glass pipette, avoiding precipitates (DO NOT STIR).
- Place in a microcentrifuge tube.
- Repeat with lead citrate.
- Centrifuge stains for 5 minutes at 13,000 rpm.

**Stain Grids with uranyl acetate**

- Using only the topmost supernatant, place drops of UA (one per grid to be stained) onto a clean Petri dish. Be sure and label positions.
- Float Grids, section side down on the surface of the droplets.
- Allow to stain for 2 minutes.
- Float grids on to a clean drop of dd $H_2O$.

**Rinse Grids thoroughly**

- One by one, pick up grids with forceps and rinse by dripping dd $H_2O$ from the squirt bottle onto the forceps above the grid. The water should then run gently over the grid.
- Holding the grid vertically over the first beaker of dd $H_2O$, dip the grid in and out of the water 30 times.
- Repeat for the other two beakers.
- Finally rinse again with squirt bottle.
- Hasten drying by blotting visible liquid with filter paper triangles between the tines of the forceps. Dry thoroughly.

**Stain with lead citrate**

- Place several pellets of NaOH into the side of the remaining Petri dish.
- Wet pellets with 1N NaOH.
- Using only the topmost supernatant, place drops of lead citrate (one per grid to be stained) onto prepared Petri dish. Label positions.
- Float Grids, section side down on the surface of the droplets.
- Allow to stain for 2 minutes.
- Float grids on to a clean drop of dd $H_2O$.

**Rinse Grids thoroughly**

- One by one, pick up grids with forceps and rinse by dripping dd $H_2O$ from the squirt bottle onto the forceps above the grid. The water should then run gently over the grid.
- Holding the grid vertically over the first beaker of dd $H_2O$, dip the grid in and out of the water 30 times.
- Repeat for the other two beakers.
- Finally rinse again with squirt bottle.
- Hasten drying by blotting visible liquid with filter paper triangles between the tines of the forceps. Dry thoroughly.
- Store in grid box.

**Clean up your area.**

**Supplemental Table 2. Grid Staining for TEM.** Protocol for using uranyl acetate and lead citrate for grid staining.
